# Aging selectively impairs how peripheral vision calibrates anticipatory postural responses to object motion

**DOI:** 10.64898/2026.05.07.723563

**Authors:** Oindrila Sinha, Isaac Kurtzer, Tarkeshwar Singh

**Affiliations:** Department of Kinesiology, The Pennsylvania State University, University Park, PA – 16802, USA; Department of Biomedical Sciences, College of Osteopathic Medicine, New York Institute of Technology, New York City, NY, USA; Neuroscience Institute, The Pennsylvania State University, University Park, PA – 16802, USA

**Keywords:** older adults, posture stabilization, smooth pursuit eye movements, peripheral vision, APA

## Abstract

Anticipatory postural adjustments (APAs) scale with velocity of approaching objects, with scaling magnitude depending on whether the moving object is actively foveated and tracked, processed through fixated peripheral vision, or processed through fixated central vision. Aging preferentially degrades the magnocellular pathway underlying peripheral motion processing while sparing the extraretinal signals available during smooth pursuit. We therefore asked whether the effect of aging on velocity-dependent APA scaling differs across these three visual pathways. Eighteen young and eighteen older adults stopped a virtual object approaching at four velocities (15–33 cm/s) under three gaze conditions: active foveation via smooth pursuit, central fixation, and peripheral fixation. We measured peak anticipatory force, rate of force development, and time to contact at force onset. Despite reduced smooth pursuit gain in older adults, velocity-dependent scaling was equivalent between age groups during active foveation, and minimal in both groups during central fixation. Critically, young adults scaled force rate during peripheral fixation nearly as steeply as during active foveation, whereas older adults’ slope was significantly lower — a difference not observed during the other gaze conditions. Older adults achieved comparable peak force by initiating responses earlier. These results establish that age-related decline in anticipatory motor control is pathway-specific: aging selectively impairs peripheral motion processing for APAs, while extraretinal mechanisms remain capable of sustaining velocity-dependent scaling. More broadly, peripheral motion processing emerges as a candidate physiological locus of age-related postural vulnerability, raising the question of whether magnocellular-targeted training can restore APA scaling in older adults.

**Key Points:**

- Young and older adults stopped virtual objects under three visual conditions: active foveation via smooth pursuit eye movements, and stationary gaze with the object moving through either central or peripheral vision.
- Velocity-dependent force rate scaling was preserved during active foveation in both age groups, minimal during fixated central vision in both age groups, and selectively impaired in older adults during fixated peripheral vision.
- We found an age-induced vulnerability in peripheral visual motion processing for anticipatory posture stabilization.

## Introduction

Everyday activities frequently require humans to interact with moving objects, such as catching a ball, or to manipulate stationary objects while the body is in motion, such as opening a door while walking. Successful completion of these tasks depends on the ability of the nervous system to anticipate and absorb mechanical forces at the moment of contact through anticipatory postural adjustments (APAs). Lacquaniti and colleagues established that APA amplitude scales linearly with object momentum (1, 2), and we have built on this work to investigate how visual motion signals are processed to modulate APAs (3, 4) and compensatory reflexes (5). The present study aimed to determine how age-related deterioration in motion processing affects the modulation of APAs in response to objects approaching at different velocities.

The most natural means of processing object motion is through smooth pursuit eye movements (SPEM), which minimize retinal image motion and provide velocity estimates via an extraretinal efference copy of the oculomotor command (6–8). Alternatively, during gaze fixation, retinal motion signals from the object’s image moving across the retina provide velocity information.

Gaze fixation leads to velocity overestimation relative to SPEM (9–11), yet we found that APA amplitudes were lower during fixation (3). We subsequently showed that this reflects two distinct sources of asymmetry: non-uniform processing across the visual field, with peripheral fixation eliciting larger APAs than central fixation, and an additional advantage for active tracking via SPEM over fixation (4).

This asymmetry reflects distinct motion-processing systems across the visual field (12–15). Central vision (∼10–15° of fovea) is dominated by a parvocellular-mediated displacement system that integrates successive stimulus positions and is optimized for low velocities below 10°/s (16–19). Peripheral vision (beyond 15°) is dominated by a magnocellular-mediated kinetic system that directly detects continuous motion across a broad velocity range (17, 18, 20, 21). Together with extraretinal efference copy signals during SPEM, these pathways constitute three distinct motion-processing mechanisms, each selectively engageable through manipulation of gaze behavior.

Aging impairs both oculomotor tracking and retinal motion processing. Older adults exhibit lower SPEM gains, resulting in poorer tracking (22–25). Critically, age-related decline is not uniform across the visual field: peripheral motion perception thresholds are elevated (26, 27), consistent with preferential deterioration of the magnocellular pathway (28–32). Despite this converging evidence, the links between age-related decline in motion processing and anticipatory posture stabilization have never been established.

Aging affects not only the visual signals that drive APAs but also APAs themselves. Older adults exhibit smaller, delayed, or temporally reorganized APAs in paradigms involving voluntary movement initiation or expected mechanical loading (33, 34) accompanied by altered cortical organization of postural control consistent with adaptive neural compensation (35). This evidence comes almost exclusively from paradigms in which the perturbation arises from voluntary movement or a force delivered to the body itself, leaving open the question of how aging affects APAs driven exclusively by visual signals from an approaching external object.

To address this gap, we asked which visual motion-processing pathway is most vulnerable to aging in driving APAs. We designed three gaze conditions that preferentially recruit each pathway: during smooth pursuit eye movements (SPEM), participants actively tracked the object, minimizing retinal image motion and relying primarily on extraretinal efference copy signals; during central fixation (FC), participants fixated a cross positioned along the object’s path so that the approaching object’s image swept across the fovea, predominantly engaging parvocellular displacement detection; and during peripheral fixation (FLR), participants fixated a cross displaced 25° laterally so that the object traversed in peripheral vision, loading most heavily on the magnocellular kinetic system. Because aging differentially affects these pathways (26), we hypothesized that the effect of aging on velocity-dependent APA scaling would be pathway-specific: impaired in conditions that depend on an age-vulnerable pathway and spared in conditions that depend on age-robust pathways. Our specific predictions were: (a) during SPEM, scaling would be preserved, because the efference copy retains proportional scaling with target velocity even as oculomotor gain declines with age; (b) during FC, both groups would show weak scaling, because the parvocellular system is poorly suited for the velocity range tested (15–33 cm/s); and (c) during FLR, older adults would show the greatest impairment, given preferential age-related degradation of the magnocellular pathway.

## Methods

### Participants

Eighteen right-hand dominant older adults (mean age: 71.2 ± 4.4 years; 9 males) and eighteen right-hand dominant younger adults (mean age: 22.2 ± 4.0 years; 9 males) participated in the study. All the participants were screened for any history of neurological conditions, visual field deficits, or musculoskeletal injuries in the upper limb. Before participating, each participant provided written informed consent and was compensated for their time ($10/h). The study protocol was approved by the Pennsylvania State University IRB (STUDY00024035) in accordance with the principles of the Declaration of Helsinki.

### Apparatus and stimuli

The experimental tasks were performed using a Kinarm Endpoint robot (Kinarm, ON, Canada) equipped with a handle force transducer. This setup was integrated with an SR EyeLink 1000 Remote eye-tracking system (SR Research, ON, Canada). Participants interacted with virtual stimuli projected from a monitor onto a semi-transparent mirror positioned above the workspace, blocking direct view of the hand. Participants held a robotic manipulandum with their right hand, with their head tilted approximately 30° forward during the experiments to facilitate tracking of eye movements. Visual stimuli, including a cursor indicating the hand’s current location, were presented at 120 Hz on a virtual reality display (VPixx Technologies, QC, Canada). Hand kinematics and kinetics were captured at 1,000 Hz. The virtual moving stimuli were generated using a Gabor-patch generator (https://www.cogsci.nl/gaborgenerator), maintaining the properties used in previous studies.

The monocular eye-tracking system was positioned roughly 80 cm from the participant’s face. Data collection was conducted in a dimly lit environment using the built-in 13-point calibration provided by KINARM. Eye movements were recorded at a maximum sampling frequency of 500 Hz with an accuracy of 0.25 – 0.5°. Slight lateral head movements were allowed, with participants resting their forehead on a pad placed on the Kinarm. The system recorded left eye movement along the x and y axes of the Kinarm’s visual surface. Total data loss was approximately 22.4 ± 14.5% (36).

### Experimental task

Participants stopped a virtual object moving at constant velocity toward them (3, 4). We simulated the physical interaction by assigning the object a virtual mass and converting the object’s momentum (mass × velocity, Eq. 1a) into a force impulse (Eq. 1a) when the virtual object contacted the cursor controlled by the hand. The impulse was delivered by the Kinarm robot as a trapezoidal force (90 ms duration; 10 ms rise/fall times, Fig. 1A). Participants were instructed to apply a counterforce within ±15% error margin to successfully stop the object (Eqs. 1b-c). After each attempt, participants received qualitative and quantitative feedback indicating their accuracy relative to the ideal ΔImpulse.

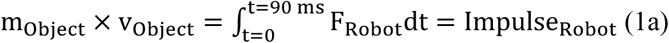

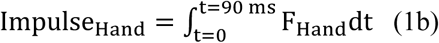

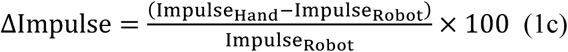

**Figure 1:**
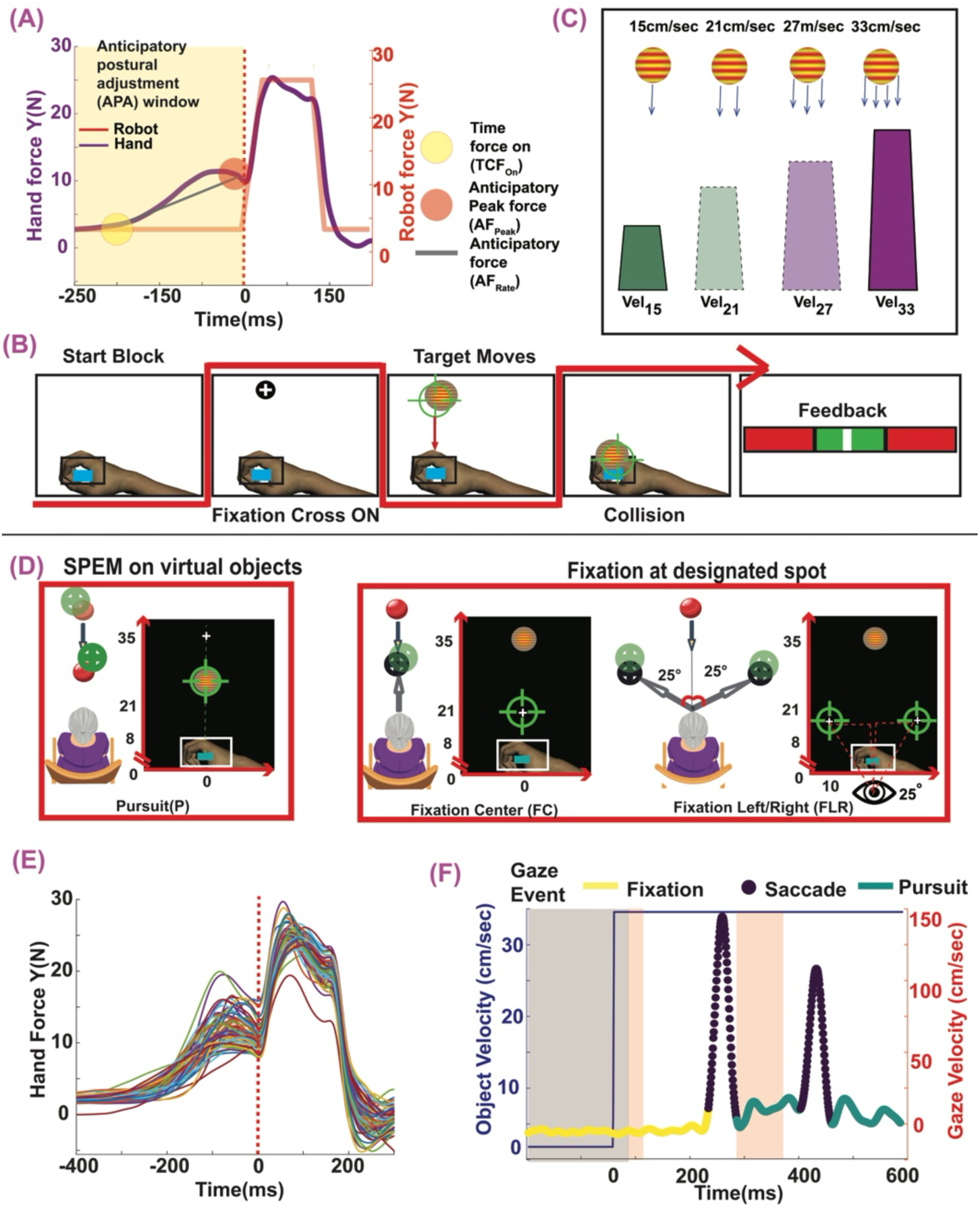
Experimental framework and task design. (**A**) Anticipatory and compensatory postural adjustments during a representative trial. Hand force (purple) and robot force (red) are shown relative to collision (time = 0). Time to contact at force onset (TCF_On_), peak anticipatory force (AF_Peak_), and rate of force development (AF_Rate_) are indicated. **(B)** Trial sequence. Participants positioned a cursor (blue rectangle) at the start position, fixated a cross, then intercepted an approaching virtual object by matching a robot-delivered force pulse. Performance feedback was provided after each trial with green areas indicating ±15% of desired force. The black rectangle indicates the area within which participants were instructed to stop the object. **(C)** Four velocity conditions (15, 21, 27, 33 cm/s) with constant object mass, producing distinct momentum values. Trapezoids indicate the force pulses that were applied for each velocity (momentum) condition. Y-axis indicates distance in cm. **(D)** Three gaze conditions: Pursuit (P), in which participants tracked the object with smooth pursuit eye movements; Fixation Centre (FC), in which gaze was held centrally while the object moved along the midline; and Fixation Left-Right (FLR), in which gaze was directed 25° laterally while the object moved in peripheral vision. **(E)** Hand force traces from one block of Vel_33_ trials (33 cm/s) for a representative participant. **(F)** Object velocity (blue, left axis) and gaze velocity (right axis) for a representative pursuit trial. Gaze events are classified as fixation (yellow), saccade (purple), or smooth pursuit (green).

At trial onset, participants positioned their hand-controlled cursor on a designated start box. A constant 3 N background force was applied throughout the trial, on top of which the impulsive force was superimposed. After cursor placement, participants fixated on a cross for 800 ms. Following fixation-cross offset, the virtual object appeared and moved toward the participant’s hand at constant velocity within 250-750 ms (Fig. 1B).

### Experimental conditions

Four speed conditions were created by varying the object’s velocity while holding mass (5.33 kg) and radius (1.5 cm) constant, resulting in distinct momentum values. Velocities were set at 15 cm/s (Vel_15_), 21 cm/s (Vel_21_), 27 cm/s (Vel_27_), and 33 cm/s (Vel_33_) (Fig. 1C).

### Study design

This study, similar to our previous work (4) was conducted over two days.

**Day 1**: Participants were familiarized with the task through a practice session using the slowest (Vel_15_) and fastest (Vel_33_) speeds in blocked conditions. For each speed, participants completed four blocks of 30 trials while tracking objects with smooth pursuit eye movements (SPEMs), referred to as the Pursuit (SPEM) condition.

**Day 2**: Participants performed 18 blocks of the MSTOP paradigm, equally divided among three randomized gaze conditions: Smooth Pursuit (SPEM, Fig. 1D, left panel), Fixation center (FC), and Fixation left-right (FLR) (Fig. 1D), with six blocks per condition. In both fixation conditions, participants fixated on a cross while the experimenter monitored compliance. The FC cross was positioned 5 cm above the starting point along the body’s midline (Fig. 1D, middle panel). In the FLR condition, the cross appeared 25° left or right of center (Fig. 1D, right panel). Each block contained 30 trials: 12 each at Vel_15_ and Vel_33_, and 3 each at Vel_21_ and Vel_27_, presented in random order. Intermediate velocities (Vel_21_, Vel_27_) made up only 10% of trials each, to probe how participants prepare APAs under less-practiced conditions

## Data recording and analyses

### Kinetic and kinematic analysis

Hand data were low-pass filtered at 50 Hz (kinetic) and 15 Hz (kinematic) using a double-pass, zero-lag, 3rd-order filter. Prior to virtual object contact, participants typically moved their hand toward the object and increased force, reaching maximum force just before contact, termed anticipatory peak force (AF_Peak_, Fig. 1A). To identify force onset, we traced force backward in time to where it fell below 5% of AF_Peak_ and identified the nearest minimum or inflection point, designated as hand force onset. We calculated the time to contact (TCF_On_) by dividing the hand-object distance by object velocity at hand force onset. A first-order polynomial fitted between the hand force at TCF_On_ and AF_Peak_ yielded the slope, quantifying the rate of force development (AF_Rate_). Example hand force data for one participant are shown in Figure 1E.

### Gaze data analyses

Gaze data were preprocessed and analyzed following established procedures (3–5). Point-of-regard (POR) data were low-pass filtered at 15 Hz, converted to spherical coordinates, and used to calculate ocular kinematics. Angular gaze speed was smoothed using a Savitzky-Golay filter (6th-order polynomial, 27-frame window). Gaze events (saccades, fixations, smooth pursuits) were identified using a threshold-based algorithm (37). Example gaze data for one trial are shown in Figure 1F.

### Statistical analyses

Three dependent variables indexing anticipatory postural adjustments were analysed: peak anticipatory force (AFPeak, N), rate of anticipatory force development (AFRate, N/ms), and time to contact (TCF_On_, ms). In addition, three gaze variables were analysed during the Pursuit condition to characterise age-related differences in smooth pursuit quality: SPEM percentage (proportion of the trial spent in smooth pursuit), gaze velocity (eye speed during pursuit, cm/s), and gaze gain (ratio of eye velocity to target velocity). All analyses were conducted in R (v 4.3) using the packages *lme4, lmerTest, emmeans*, and *afex*. The significance level was set at α = 0.05 throughout.

#### Age group comparison

To establish whether overall motor performance differed between age groups, an independent-samples Welch’s t-test compared grand-mean TCF_On_, AF_Rate_, and AF_Peak_ (collapsed across all gaze and speed conditions) between young and older adults.

#### Gaze variable analyses

To characterize age-related differences in smooth pursuit eye movements, a two-way mixed ANOVA was conducted for each gaze variable (SPEM percentage, gaze velocity, gaze gain) on data from the Pursuit condition only, with age group (Young, Older) as a between-subjects factor and speed condition (Vel_15_, Vel_21_, Vel_27_, Vel_33_) as a within-subjects factor. Post-hoc independent-samples t-tests compared age groups at each speed level with Holm–Bonferroni correction.

#### Linear mixed-effects models

We employed a linear mixed-effects model (LMM) treating target velocity as a continuous predictor (38). The LMM estimates the slope, how steeply each motor variable scales per unit increase in target velocity (1 df), and tests whether these slopes differ across gaze conditions and age groups via interaction terms.

For each dependent variable, a single LMM was fit to all trial-level data (15,085 trials from 36 participants) using restricted maximum likelihood estimation:

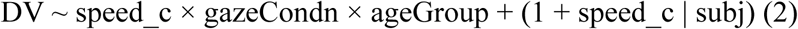

where speed_c is target velocity mean-centered at 24 cm/s (the grand mean of 15, 21, 27, and 33 cm/s), gazeCondn is a three-level factor (Fixation Centre [FC], Fixation Left-Right [FLR], and Pursuit [P]), and ageGroup is a two-level between-subjects factor (Young and Older). The random-effects structure included a by-subject random intercept and a by-subject random slope for speed_c, allowing each participant to have their own baseline motor output and their own sensitivity to target velocity. This structure accounts for the repeated-measures dependency among trials within participants and ensures that the effective sample size for the between-subjects factor (age group) is based on the number of participants rather than the number of trials. Significance of fixed effects was evaluated using Type III F-tests with Satterthwaite degrees of freedom.

The key fixed-effect terms map directly onto our hypotheses. The coefficient for speed_c estimates the overall slope, or how much the dependent variable changes per 1 cm/s increase in target velocity. The speed_c × gazeCondn interaction tests whether this slope differs across gaze conditions. The gazeCondn × ageGroup interaction tests whether the two age groups differ in mean motor output across gaze conditions. Critically, the three-way speed_c × gazeCondn × ageGroup interaction tests whether the two age groups modulate their velocity-dependent scaling (slopes) differently across gaze conditions — the central test of our hypothesis that aging differentially impairs visuomotor gain across gaze conditions that preferentially engage distinct motion-processing pathways.

#### Post-hoc decomposition

Following significant interactions, estimated marginal means (emmeans) were computed to compare mean motor output across conditions and groups at the average target velocity. Estimated marginal trends (emtrends) were computed to extract the slope of each dependent variable on speed_c for each gaze condition × age group combination. Two post-hoc paths were followed: (a) pairwise comparisons of slopes across gaze conditions within each age group, testing whether velocity-dependent scaling differs across gaze conditions; and (b) pairwise comparisons of slopes between age groups within each gaze condition, testing whether aging impairs velocity-dependent scaling at a given gaze condition. All post-hoc comparisons were adjusted for multiple comparisons using the Holm–Bonferroni method. Descriptive statistics are reported as mean ± SD unless otherwise noted as SE.

#### Linearity check

Because the LMM treats target velocity as a continuous linear predictor, we tested whether a quadratic component improved fit. For each dependent variable, we refit the LMM with an additional fixed-effect term I(speed_c²) and compared it to the linear model via a likelihood-ratio test (ML estimation, 1 df). The quadratic term did not significantly improve fit for TCF_On_ (χ²(1) = 1.30, p = .255) or AF_Peak_ (χ²(1) = 1.55, p = .213). For AF_Rate_ the improvement was marginal (χ²(1) = 3.32, p = .07); however, the estimated quadratic coefficient was 1.4 × 10⁻⁵, contributing at most ∼13% of the linear effect over the velocity range tested, and was therefore considered negligible. We retain the linear specification for all subsequent analyses.

## Results

### Older adults exhibit reduced smooth pursuit gain despite comparable pursuit engagement

To verify that the expected age-related decline in oculomotor function was present in our sample, we examined three gaze variables during the Pursuit condition: the proportion of trial time spent in smooth pursuit (SPEM percentage), gaze velocity, and gaze gain (Figure 2).

**Figure 2:**
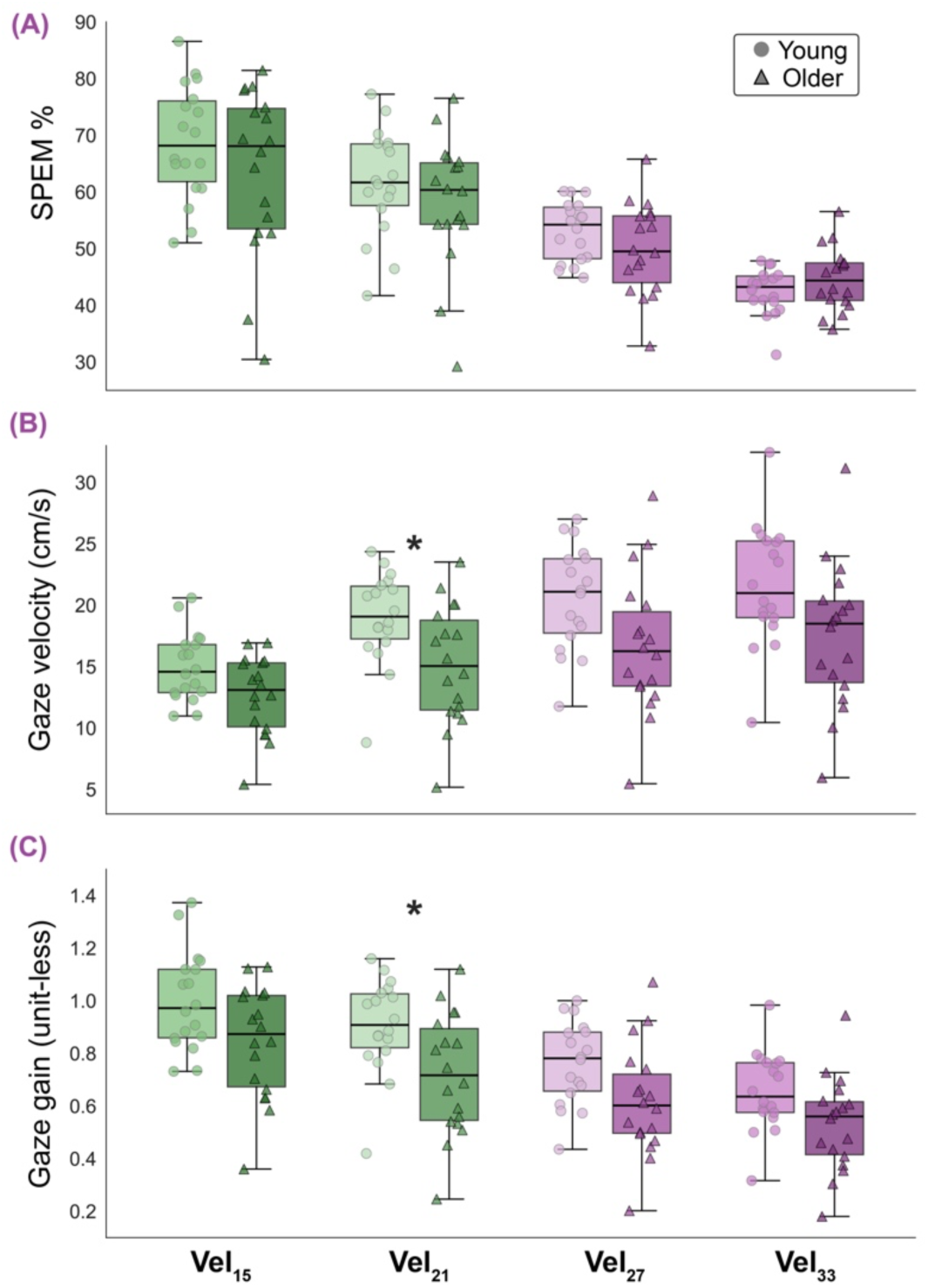
Older adults exhibit reduced smooth pursuit gain. Three gaze variables measured during the Pursuit condition are shown as a function of target velocity for young (circles) and older (triangles) adults. (A) SPEM percentage: proportion of trial time spent in smooth pursuit eye movements. Both groups showed equivalent pursuit engagement (p = .34), with SPEM percentage declining with faster targets (p < .001). (B) Gaze velocity: eye speed during pursuit. Older adults tracked significantly more slowly than young adults (p = .016). (C) Gaze gain: ratio of eye velocity to target velocity. Older adults had significantly lower gain (0.68 vs 0.83, p = .016), indicating their eyes lagged further behind the target. No age × speed interactions were significant for any variable. Boxplots show medians and interquartile ranges; individual data points represent subject-level means at each speed. Asterisks denote significant age group differences after Holm–Bonferroni correction (*p < .05).

Young and older adults spent a comparable proportion of each trial engaged in smooth pursuit (Young: 56.5 ± 12.4%; Old: 54.2 ± 12.6%; *F*(1, 34) = 0.95, *p* = .34, η²_p_ = .027, Fig. 2A). SPEM percentage declined sharply with increasing target velocity (*F*(1.56, 53.16) = 80.53, *p* < .001, η²p = .70), dropping from approximately 66% at Vel_15_ (15 cm/s) to 44% at Vel_33_ (33 cm/s) as faster targets became progressively harder to track smoothly. The age × speed interaction was not significant (*F*(1.56, 53.16) = 1.98, *p* = .16), indicating that both groups experienced the same decline in pursuit engagement with increasing target velocity. Thus, older adults did not compensate for tracking difficulty by switching to a saccadic strategy more than younger adults.

Despite spending equivalent time in smooth pursuit, older adults tracked the target with significantly lower gaze velocity (Young: 19.1 ± 4.7 cm/s; Old: 15.5 ± 5.2 cm/s; *F*(1, 34) = 6.37, *p* = .016, η²_p_ = .16, Fig. 2B) and lower gaze gain (Young: 0.83 ± 0.21; Old: 0.68 ± 0.23; *F*(1, 34) = 6.49, *p* = .016, η²_p_ = .16, Fig. 2C). Both measures increased with target velocity (gaze velocity: *F*(2.16, 73.56) = 69.19, *p* < .001, η²p = .67; gaze gain: *F*(2.57, 87.32) = 159.24, *p* < .001, η²p = .82), but the age × speed interactions were not significant (gaze velocity: *p* = .15; gaze gain: *p* = .32), confirming that the age-related reduction in pursuit quality was uniform across target velocities. Post-hoc comparisons between age groups at each speed level showed consistent trends at all four velocities (all uncorrected *p* < .03), though only the comparison at Vel_21_ (21 cm/s) survived Holm–Bonferroni correction (*p* = .043) for both gaze velocity and gaze gain.

In summary, older adults maintained the same pursuit pattern as younger adults: they did not reduce the proportion of time spent tracking, but their eyes moved more slowly relative to the target. The reduced gaze velocity and gain imply greater residual retinal image motion during pursuit in older adults, which could affect the quality of the velocity estimate available for calibrating anticipatory motor responses.

### Older adults initiate force earlier during pursuit and peripheral fixation but not central fixation

Young and older adults did not differ in overall time to contact (Young: −287 ± 79 ms; Old: −320 ± 83 ms; *t*(33.9) = 1.22, *p* = .23). Both groups initiated force well before collision, and the magnitude of this temporal margin was comparable.

The LMM revealed significant main effects of speed (*F*(1, 18.9) = 113.96, *p* < .001) and gaze condition (*F*(2, 15040) = 20.08, *p* < .001), with no overall age difference (*p* = .33). The speed × gaze interaction was significant (*F*(2, 14997) = 7.49, *p* < .001), confirming that the velocity–timing relationship differed across gaze conditions. The gaze × age interaction was significant (*F*(2, 15040) = 48.08, *p* < .001), indicating that the two age groups distributed their timing differently across gaze conditions. The three-way speed × gaze × age interaction was marginal (*F*(2, 14997) = 2.82, *p* = .06).

Estimated marginal means revealed a clear dissociation between groups (Fig. 3A). Young adults showed only small differences in TCF_On_ across gaze conditions (range of means ∼12 ms; FC: −291 ms; FLR: −288 ms; P: −279 ms), with active pursuit associated with slightly later force initiation than either fixation condition (FC vs Pursuit, p < .001; FLR vs Pursuit, p = .001; FC vs FLR not significant, p = .33). In contrast, older adults initiated force substantially earlier during Pursuit (−328 ms) and FLR (−331 ms) than during central fixation (FC: −302 ms; both p < .001), while Pursuit and FLR did not differ from each other (p = .30), yielding a wider range of means (∼29 ms) than in young adults. This reorganization was confirmed by a significant gaze × age interaction (F(2,15040) = 48.08, p < .001): peripheral fixation clustered with central fixation in young adults but with pursuit in older adults. Despite these within-group differences in pattern, direct age comparisons within each gaze condition were not significant (all p > .05).

**Figure 3:**
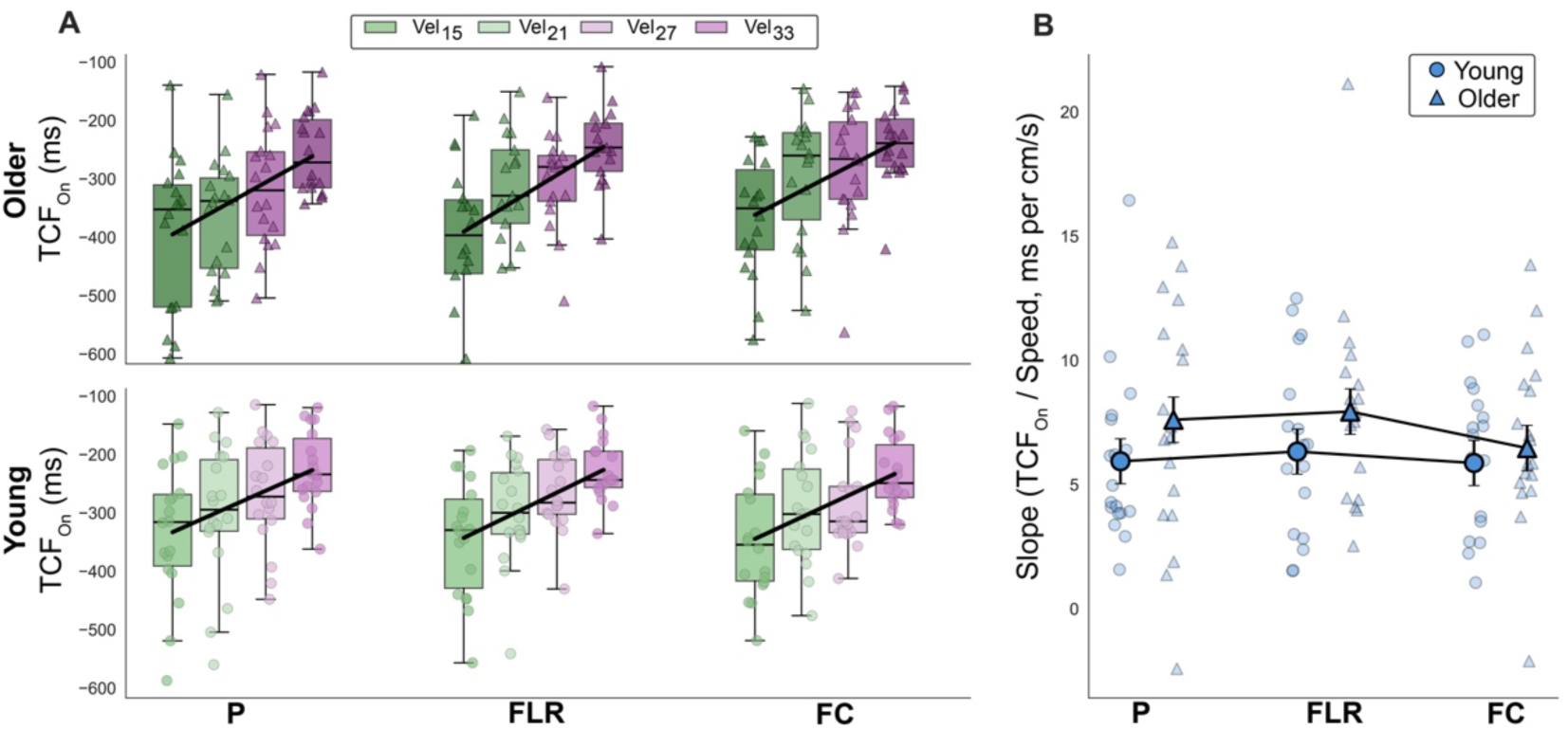
Older adults initiate force earlier during pursuit and peripheral fixation but not central fixation. (**A**) Time to contact at force onset (TCF_On_) across gaze conditions (P: Pursuit; FLR: Fixation Left-Right; FC: Fixation Centre) and target velocities (15–33 cm/s) for older adults (top row, triangles) and young adults (bottom row, circles). Boxplots show subject-level means at each speed; colors grade from green (15 cm/s) to purple (33 cm/s). Black lines show group-mean OLS regression slopes of TCF_On_ on target velocity. More negative values indicate earlier force initiation. Young adults’ timing was gaze-invariant, whereas older adults initiated force ∼28 ms earlier during Pursuit and FLR than during FC. (B) Velocity-dependent slopes of TCF_On_ (ms per cm/s) estimated from the linear mixed-effects model (emtrends). Large symbols with error bars show LMM-estimated group slopes ± SE; small semi-transparent symbols show individual participant OLS slopes. Young adults’ slopes were uniform across gaze conditions (∼6 ms per cm/s). Older adults showed steeper slopes during Pursuit (P) and FLR than FC (both p < .01). No direct age comparisons of slopes were significant at any gaze condition.

Estimated slopes (emtrends) showed that young adults scaled their TCFOn uniformly with target velocity across all gaze conditions (FC: 5.87; FLR: 6.33; P: 5.94 ms per cm/s; all pairwise *p* > .52; Fig. 3B). Older adults, by contrast, showed steeper slopes during FLR (7.94 ms per cm/s) and Pursuit (7.61 ms per cm/s) than during FC (6.47 ms per cm/s; FC vs FLR: *p* < .001; FC vs P: *p* = .005), while FLR and Pursuit did not differ (*p* = .382; Fig. 3B). The steeper slopes indicate that older adults’ earlier force initiation during pursuit and peripheral fixation became more pronounced at higher target velocities. Direct age comparisons of slopes within each gaze condition were not significant (all *p* > .19).

In summary, young adults’ force initiation timing was largely insensitive to gaze condition, whereas older adults initiated force earlier and with steeper velocity-dependent scaling during pursuit and peripheral fixation compared to central fixation, where both timing and its velocity dependence were attenuated.

### Velocity-dependent scaling of anticipatory force rate is selectively impaired during peripheral fixation in older adults

Overall anticipatory force rate (Young: 0.04 ± 0.016 N/ms; Old: 0.038 ± 0.017 N/ms; *t*(33.9) = 0.392, *p* = .7) and peak anticipatory force (Young: 7.34 ± 0.98 N; Old: 7.35 ± 1.47 N; *t*(29.6) = −0.011, *p* = .99) did not differ between age groups. The two groups therefore produced comparable overall force output despite the oculomotor differences documented above.

### Anticipatory force rate (AF_Rate_)

The LMM revealed significant main effects of speed (*F*(1, 33.8) = 74.68, *p* < .001) and gaze condition (*F*(2, 15027) = 306.79, *p* < .001), with no overall age difference (*p* = .68). The speed × gaze (*F*(2, 15030) = 133.98, *p* < .001) and gaze × age (*F*(2, 15027) = 50.46, *p* < .001) interactions were both significant. Critically, the three-way speed × gaze × age interaction was significant (*F*(2, 15030) = 6.99, *p* < .001), indicating that the two age groups modulated their velocity-dependent force rate scaling differently across gaze conditions.

Estimated marginal means revealed a clear hierarchy in young adults: FC (0.034 N/ms) < FLR (0.040 N/ms) < Pursuit (0.046 N/ms), with all pairwise comparisons significant (all *p* < .001; Fig. 4A). In older adults, this hierarchy was disrupted: FC (0.034 N/ms) was significantly higher than FLR (0.032 N/ms; *p* = .007), and both were substantially lower than Pursuit (0.048 N/ms; both *p* < .001). The reversal of the FC–FLR ordering — young adults showed higher force rates during peripheral fixation, whereas older adults did not — contributed to the significant three-way interaction. Direct age comparisons within each gaze condition were not significant (all *p* >.11).

**Figure 4:**
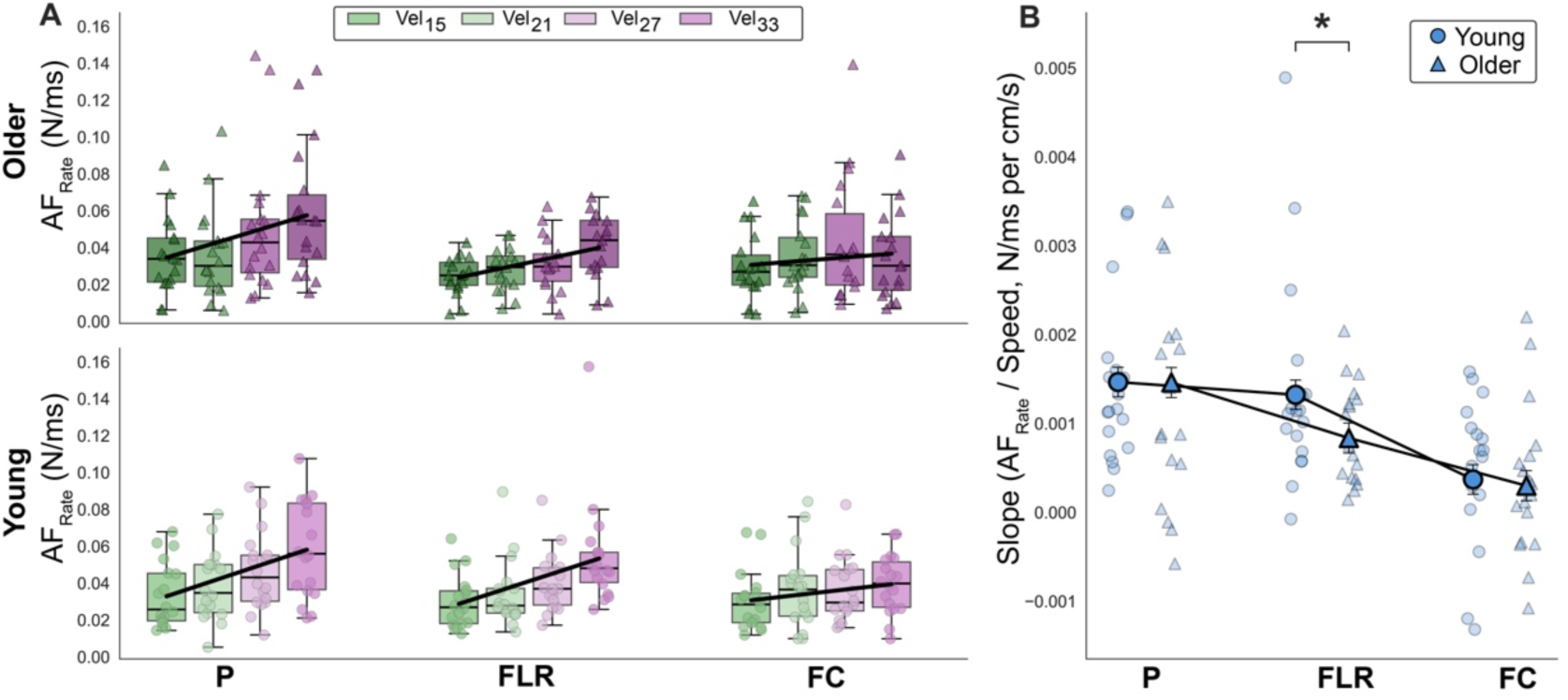
Velocity-dependent scaling of anticipatory force rate is selectively impaired during peripheral fixation in older adults. (**A**) Anticipatory force rate (AF_Rate_, N/ms) across gaze conditions and target velocities for older adults (top row, triangles) and young adults (bottom row, circles). Format as in Fig. 3A. Both groups showed significant gaze × speed interactions (Young: p = .003; Old: p = .008). Young adults exhibited a clear hierarchy: FC < FLR < Pursuit (P). In older adults, this hierarchy was disrupted: FLR did not exceed FC. (B) Velocity-dependent slopes of AF_Rate_ (N/ms per cm/s) from the LMM. Format as in Fig. 3B. The three-way speed × gaze × age interaction was significant (p < .001). In young adults, the FLR slope was close to Pursuit (p = .13); in older adults, the FLR slope dropped approximately halfway between FC and Pursuit (all pairwise p < .001). The asterisk denotes a significant age difference in slope at FLR (p = .04); no age differences were observed at FC (p = .77) or Pursuit (p = .98).

Estimated slopes revealed that both groups showed the same general hierarchy — Pursuit > FLR > FC — but differed in the spacing between conditions (Fig. 4B). In young adults, the FLR slope (0.00132 N/ms per cm/s) was close to Pursuit (0.00147; FLR vs P: *p* = .13), and both were significantly steeper than FC (0.00037; both *p* < .001). In older adults, the FLR slope (0.00084) fell between FC (0.0003) and Pursuit (0.00146), with all three pairwise comparisons significant (all *p* < .001). Whereas young adults’ peripheral fixation yielded force rate scaling nearly equivalent to active pursuit tracking, older adults’ FLR slope dropped approximately halfway between FC and Pursuit, indicating that aging substantially reduced the benefit that peripheral vision provides for velocity-dependent force rate modulation in young adults. Direct age comparisons of slopes confirmed that the groups did not differ at FC (*p* = .768) or Pursuit (*p* = .98), but older adults had a significantly flatter slope at FLR (*p* = .04). Thus, both groups showed minimal velocity-dependent scaling during central fixation, whereas the absence of an age difference during Pursuit indicates that velocity estimation adequate for scaling anticipatory force is preserved with aging through extraretinal mechanisms. The selective age-related reduction at FLR is consistent with deterioration of the magnocellular pathway that subserves peripheral motion processing.

### Peak anticipatory force (AF_Peak_)

The LMM confirmed significant main effects of speed (*F*(1, 33) = 114.82, *p* < .001) and gaze (*F*(2, 15022) = 640.53, *p* < .001), with no age difference (*p* = .98). The speed × gaze (*F*(2, 15030) = 223.27, *p* < .001) and gaze × age (*F*(2, 15022) = 71.67, *p* < .001) interactions were significant. However, the three-way interaction was not significant (*F*(2, 15030) = 0.41, *p* = .67), indicating that both groups modulated their peak force scaling identically across gaze conditions.

Estimated marginal means showed the same Pursuit > FLR > FC hierarchy in both groups, with all pairwise comparisons significant (all *p* < .001; Fig. 5A). Older adults exhibited a larger Pursuit–FC separation (1.51 N) than young adults (0.78 N), driving the gaze × age interaction. Direct age comparisons within each gaze condition were not significant (all *p* > .30).

**Figure 5:**
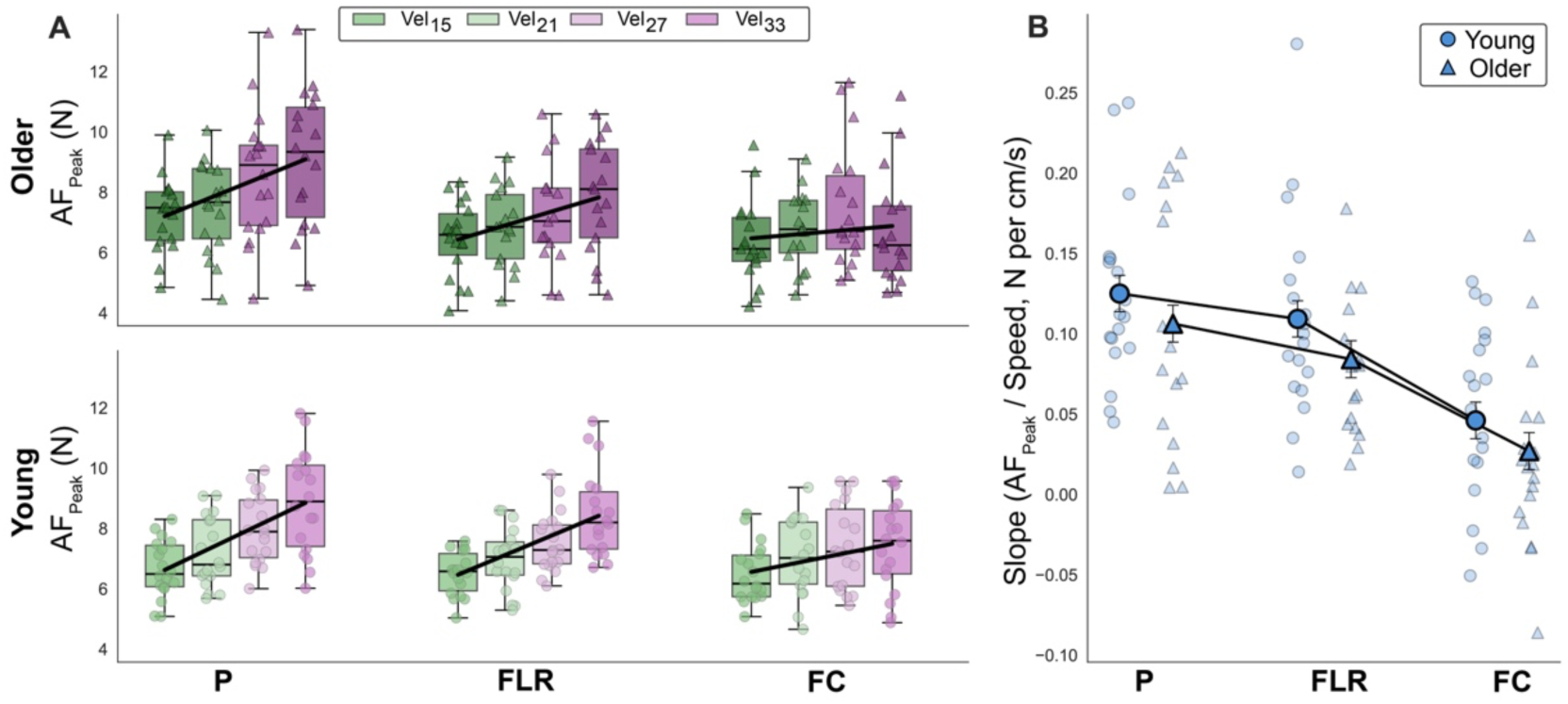
Velocity-dependent scaling of peak anticipatory force is preserved across all gaze conditions in both age groups. (**A**) Peak anticipatory force (AF_Peak_, N) across gaze conditions and target velocities for older adults (top row, triangles) and young adults (bottom row, circles). Format as in Fig. 3A. All three effects (gaze, speed, gaze × speed) were significant in both groups. Both groups showed the same Pursuit (P) > FLR > FC hierarchy. Older adults exhibited a larger Pursuit–FC separation (1.51 N) than young adults (0.78 N). (B) Velocity-dependent slopes of AF_Peak_ (N per cm/s) from the LMM. Format as in Fig. 3B. The three-way speed × gaze × age interaction was not significant (p = .67). Both groups showed identical slope hierarchies (Pursuit > FLR > FC) with all pairwise comparisons significant (all p < .01). No age differences in slope were observed at any gaze condition (all p > .12), confirming that velocity-dependent scaling of peak force was fully preserved in older adults despite the selective impairment of force rate scaling during peripheral fixation.

Estimated slopes confirmed that both groups scaled peak force with target velocity in an identical pattern: Pursuit > FLR > FC, with all pairwise comparisons significant in both groups (all *p* < .01; Fig. 5B). Young adults’ slopes were FC: 0.046, FLR: 0.109, Pursuit: 0.125 N per cm/s; older adults’ slopes were FC: 0.027, FLR: 0.084, Pursuit: 0.106 N per cm/s. Direct age comparisons of slopes were not significant at any gaze condition (all *p* > .12), confirming that the velocity-dependent scaling of peak force was fully preserved in older adults.

### Dissociation between force dynamics and force amplitude

The contrast between AF_Rate_ and AF_Peak_ is notable. The significant three-way interaction for AF_Rate_ (*p* < .001) but not AF_Peak_ (*p* = .67) reveals that aging selectively affects how quickly force is built up — not how much force is ultimately produced. For AF_Rate_, the age-related impairment is specific to peripheral fixation: older adults retain normal velocity-dependent scaling during both central fixation (where scaling is minimal in both groups) and pursuit tracking (where extraretinal signals compensate), but lose the peripheral vision advantage that young adults exploit to achieve pursuit-like force rate modulation. Older adults compensate for their slower force rate by initiating force earlier (see TCF_On_ results), ultimately reaching comparable peak force amplitudes across all gaze conditions.

### Pursuit tracking quality predicts force rate scaling in older adults

To test whether pursuit tracking quality improves force calibration beyond object velocity alone, we fit nested regression models (with and without gaze velocity as a predictor) to subject×speed means from the Pursuit condition (72 observations per group). The improvement in variance explained (ΔR²) was evaluated using a permutation test (10,000 iterations).

For peak anticipatory force, gaze velocity significantly improved prediction in both young (ΔR² = .085, p = .005) and older adults (ΔR² = .103, p = .006; Fig. 6). For force rate, the pattern diverged: gaze velocity did not significantly improve prediction in young adults (ΔR² = .019, p = .238) but explained a substantial additional 17.6% of variance in older adults (ΔR² = .176, p < .001; Fig. 6). This dissociation indicates that young adults’ force dynamics are relatively robust to variation in pursuit quality, whereas older adults’ rate of force development is tightly coupled to how well they track the approaching object.

**Figure 6:**
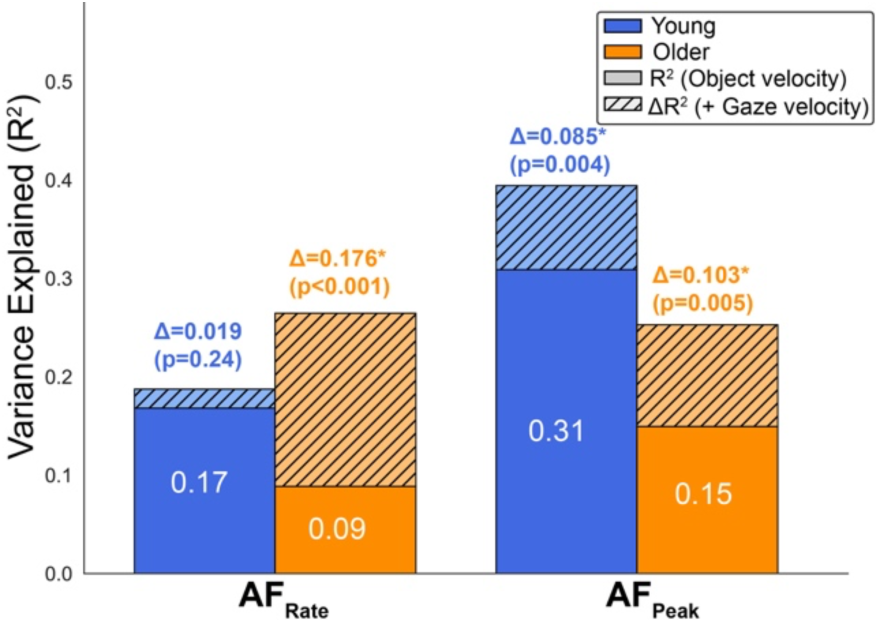
Pursuit tracking quality improves force calibration beyond object velocity alone, with older adults benefiting disproportionately for force rate. Stacked bars show variance explained (R²) in anticipatory force rate (AF_Rate_, left) and peak anticipatory force (AF_Peak_, right) for young (blue) and older (orange) adults during the Pursuit condition. Solid bars represent R² from a regression model with object velocity as the sole predictor. Hatched bars represent the additional variance explained (ΔR²) when gaze velocity is added as a predictor. Data were averaged across trials within each subject and speed condition (72 observations per group: 18 subjects × 4 target velocities). Significance of ΔR² was assessed using a permutation test (10,000 iterations) in which gaze velocity was shuffled across subjects within each speed condition. Asterisks and p-values are shown above each bar. For AF_Peak_, gaze velocity significantly improved prediction in both age groups. For AF_Rate_, gaze velocity significantly improved prediction only in older adults, indicating that their force dynamics are more dependent on the quality of their pursuit tracking signal.

## Discussion

Our central hypothesis was that the effect of aging on velocity-dependent APA scaling would be pathway-specific: impaired in conditions that depend on an age-vulnerable pathway and spared in conditions that depend on age-robust pathways. The three gaze conditions provided a direct test of these predictions.

*Prediction (a): preserved scaling during SPEM.* We predicted that scaling would be preserved because the efference copy retains proportional velocity information even as oculomotor gain declines with age. This was borne out: slopes at Pursuit did not differ between age groups for any motor variable (all p > .19), and both groups achieved their steepest velocity-dependent scaling during pursuit despite reduced gaze gain in older adults (0.68 vs 0.83). Greater residual retinal slip in older adults may additionally have provided a supplementary compensatory velocity signal.

*Prediction (b): weak scaling during central fixation in both groups*. We predicted weak scaling in both age groups because the parvocellular displacement system is poorly suited to the velocity range tested (15–33 cm/s), a limitation that is independent of age (18). As predicted, both groups showed minimal scaling at FC (e.g., AF_Rate_ slopes of 0.00037 and 0.0003 N/ms per cm/s), with no age differences at FC for any variable (all p > .24).

*Prediction (c): impaired scaling during peripheral fixation in older adults.* We predicted the greatest age-related impairment here, given preferential magnocellular decline. This held for the rate of force development: the three-way speed × gaze × age interaction was significant for AF_Rate_ (p < .001), and the direct age comparison of slopes at FLR confirmed that older adults had significantly flatter velocity-dependent force rate scaling than young adults (p = .04). In young adults, peripheral fixation yielded force rate modulation nearly equivalent to pursuit; in older adults, this peripheral advantage was substantially diminished. This selective impairment at FLR is consistent with age-related degradation of the magnocellular pathway and dorsal stream processing that subserve peripheral motion perception. However, the impairment was confined to force dynamics: the three-way interaction for AF_Peak_ was not significant (p = .67), and peak force scaling did not differ between groups at any gaze condition. For TCF_On_, the three-way interaction was marginal (p = .06), with a suggestive but non-significant pattern in the same direction.

*Compensatory strategy.* Although older adults showed impaired force rate scaling during peripheral fixation, they achieved comparable peak force by initiating force earlier. The TCF_On_ results revealed that older adults began their anticipatory response approximately 28 ms earlier during pursuit and peripheral fixation relative to central fixation, a pattern absent in young adults. This earlier initiation effectively compensated for the slower rate of force build-up, allowing older adults to reach the same peak force amplitudes as young adults across all gaze conditions. The dissociation between AF_Rate_ (significant three-way interaction) and AF_Peak_ (non-significant three-way interaction) thus reflects a shift in the temporal dynamics of anticipatory control rather than a deficit in its final output (39).

### Age-related impairment of anticipatory motor scaling is specific to the magnocellular motion-processing pathway

Our overarching hypothesis — that the effect of aging on velocity-dependent APA scaling would be pathway-specific rather than uniform — was supported by a consistent pattern across three motor variables. The three gaze conditions, each designed to selectively engage a distinct motion-processing mechanism, revealed a clear dissociation: pursuit-based APA scaling was equivalent between groups, central-fixation-based scaling was weak in both groups, and peripheral-fixation-based scaling was selectively reduced in older adults.

The preservation of APA scaling during pursuit tracking was observed despite a significant age-related reduction in gaze gain (0.68 vs 0.83, p = .016). This finding suggests that the extraretinal efference copy signal retains its proportional modulation with target velocity even when the oculomotor command itself is weaker. One explanation is that the efference copy encodes the intended eye velocity command rather than the achieved eye velocity (40–42), and the scaling of this command with target velocity is preserved even though its absolute magnitude is reduced. An alternative and complementary explanation is that the reduced gaze gain in older adults produces greater residual retinal image motion during pursuit, and this retinal slip provides a supplementary velocity signal that compensates for any degradation in the efference copy.

During pursuit, velocity information is therefore available from two partially redundant sources — extraretinal and retinal — and this redundancy may buffer the system against the effects of aging on either source alone. This interpretation is consistent with evidence that motor performance during interception tasks is most accurate when both retinal and extraretinal signals are available (8).

The minimal scaling at FC is consistent with parvocellular displacement detection. Although the mapping between parvocellular/magnocellular pathways and central/peripheral vision is not absolute (16, 17, 21), the predominance of magnocellular projections in peripheral retina and the disproportionate age-related decline in peripheral motion perception (26, 27, 43) support a functionally meaningful distinction. Our conditions therefore preferentially engage, rather than purely isolate, each pathway.

The near-zero slopes at central fixation (FC) confirm that neither group could effectively extract graded velocity information through the parvocellular pathway at the speeds tested, extending our previous findings in young adults (3, 4) to older adults and demonstrating that this limitation is intrinsic to the pathway. Although aging weakens suppression of task-irrelevant central motion signals (26), this would not improve velocity extraction for APA calibration.

The selective impairment of APA scaling during peripheral fixation in older adults represents the central finding of this study. In young adults, peripheral fixation yielded velocity-dependent force rate modulation nearly equivalent to pursuit, consistent with our previous demonstration that the magnocellular kinetic system likely provides motion signals of sufficient quality to support precise APA calibration in healthy young adults (4). In older adults, however, the FLR slope dropped approximately halfway between FC and Pursuit, and the direct age comparison at FLR was significant for AF_Rate_ (p = .04) — the only gaze condition at which slopes differed between groups. This pattern is consistent with converging evidence that aging preferentially degrades the magnocellular pathway and the dorsal visual stream that it feeds (29, 44, 45). Psychophysical studies have demonstrated elevated motion perception thresholds in the peripheral visual field of older adults (28, 31, 46), reduced contrast sensitivity in magnocellular-mediated channels (47, 48), and impaired peripheral motion detection relevant to hazard perception in real-world tasks (27). Our results provide the first direct evidence that this age-related decline in peripheral motion processing translates into impaired anticipatory postural control — specifically, a reduced capacity to scale the rate of force development with object velocity when motion must be processed primarily through the magnocellular system.

### Older adults compensate for slower force dynamics by initiating anticipatory responses earlier, preserving force amplitude

A striking feature of the present results is the dissociation between force dynamics and force amplitude. The three-way speed × gaze × age interaction was significant for AF_Rate_ (p < .001) but not for AF_Peak_ (p = .67), indicating that aging affects how quickly anticipatory force is built up without altering how much force is ultimately produced. This dissociation raises the question of how older adults achieve comparable peak force despite a slower rate of force development during peripheral fixation.

The TCF_On_ results suggest a compensatory mechanism: older adults initiated force ∼28 ms earlier during pursuit and peripheral fixation relative to central fixation, a pattern absent in young adults. By extending the temporal window for force accumulation, older adults compensated for the reduced rate, achieving comparable peak force across all conditions.

This pattern parallels findings from precision grip studies. Cole and colleagues demonstrated that age-related differences in fingertip force control during lifting tasks reflect deteriorating capacity to rapidly encode surface properties and adjust grip forces online, rather than a fundamental deficit in the ability to generate anticipatory force programs (39). In their paradigm, older adults produced comparable anticipatory grip forces to young adults but were slower to modulate these forces in response to trial-by-trial changes in surface friction. Our results extend this principle from precision grip to whole-limb anticipatory postural control during object interception: the preparatory, learned component of the response (peak force amplitude scaled to object velocity) is preserved, while the dynamic, sensory-driven component (the rate at which force is modulated in response to real-time velocity signals from peripheral vision) is impaired.

This compensatory strategy, however, may be fragile outside the laboratory. The MSTOP paradigm afforded predictable trajectories, constant velocities, and sufficient viewing time — conditions that allowed older adults to exploit the temporal margin. In everyday situations involving unexpected collisions or objects appearing from the periphery without advance warning (49), the temporal buffer may be insufficient, leaving older adults unable to generate adequate postural stabilization before contact.

### Impaired peripheral motion processing as a contributor to collision-related falls in older adults

Falls are the leading cause of injury-related death in adults over 65, with 30–50% involving collisions with environmental objects (50–53). Unlike spontaneous balance failures, collision-induced destabilization arises when the visual system fails to estimate an approaching object’s properties in time to prepare appropriate postural stabilization — yet the underlying visual processing mechanisms have remained largely unexplored.

We demonstrated that aging selectively impairs velocity-dependent scaling of anticipatory force rate during peripheral fixation — the condition that engages the magnocellular pathway most heavily. In everyday life, objects approaching from the side or from outside the current focus of attention are initially detected through peripheral vision, precisely the pathway we have shown to be compromised in older adults. If the magnocellular system cannot provide accurate velocity signals for these peripherally detected objects, the motor system will be unable to calibrate anticipatory postural responses appropriately, increasing the likelihood that contact with the object will result in postural destabilization. This interpretation is supported by recent work demonstrating that age-related decline in peripheral motion perception is associated with poorer performance on hazard perception tasks that require detecting and responding to objects approaching in the visual periphery (27). As noted above, the compensatory strategy of earlier force initiation observed in our paradigm depends on predictable trajectories and sufficient viewing time and may be insufficient under these real-world conditions.

Older adults’ rate of force development was tightly coupled to their pursuit quality, whereas young adults’ was not (Fig. 6) — a dissociation that, combined with the preserved velocity-dependent scaling during pursuit, suggests that actively foveating approaching objects via smooth pursuit eye movements could serve as a viable compensatory strategy for maintaining adequate anticipatory postural control in aging.

These findings point toward two potential intervention strategies. First, training programs that encourage active visual tracking of moving objects, rather than passive peripheral detection, could help older adults maintain accurate velocity estimation and appropriate anticipatory postural control. Second, interventions targeting rapid force production, such as power training, may help older adults generate adequate postural responses within shorter temporal windows when compensatory early initiation is not possible. Future work should examine whether the peripheral processing deficit identified here generalizes to more naturalistic collision scenarios involving unpredictable approach trajectories, and whether targeted visual or motor training can mitigate the age-related decline in magnocellular-mediated anticipatory postural control.

### A unifying conceptual model and predictions

The findings described above invite a more systematic account of how aging reshapes the computations that links visual motion to anticipatory motor output. Figure 7 presents a signal-flow architecture in which three motion-processing pathways converge on a velocity estimate *v*” characterized by both magnitude and precision κ; magnitude drives force-rate generation, while precision drives a parallel timing computation that shifts force onset earlier when reliability is reduced. Three results constrain the model. The preservation of pursuit-based APA scaling with age suggests that the extraretinal coupling g_E_ is unaffected. The selective deficit at peripheral fixation points to the magnocellular coupling g_M_ as the site of impairment. And the fact that older adults shift force onset earlier during SPEM and peripheral fixation but not during central fixation suggests that this temporal compensation is adopted only when the velocity estimate eventually becomes reliable enough to act upon.

**Figure 7:**
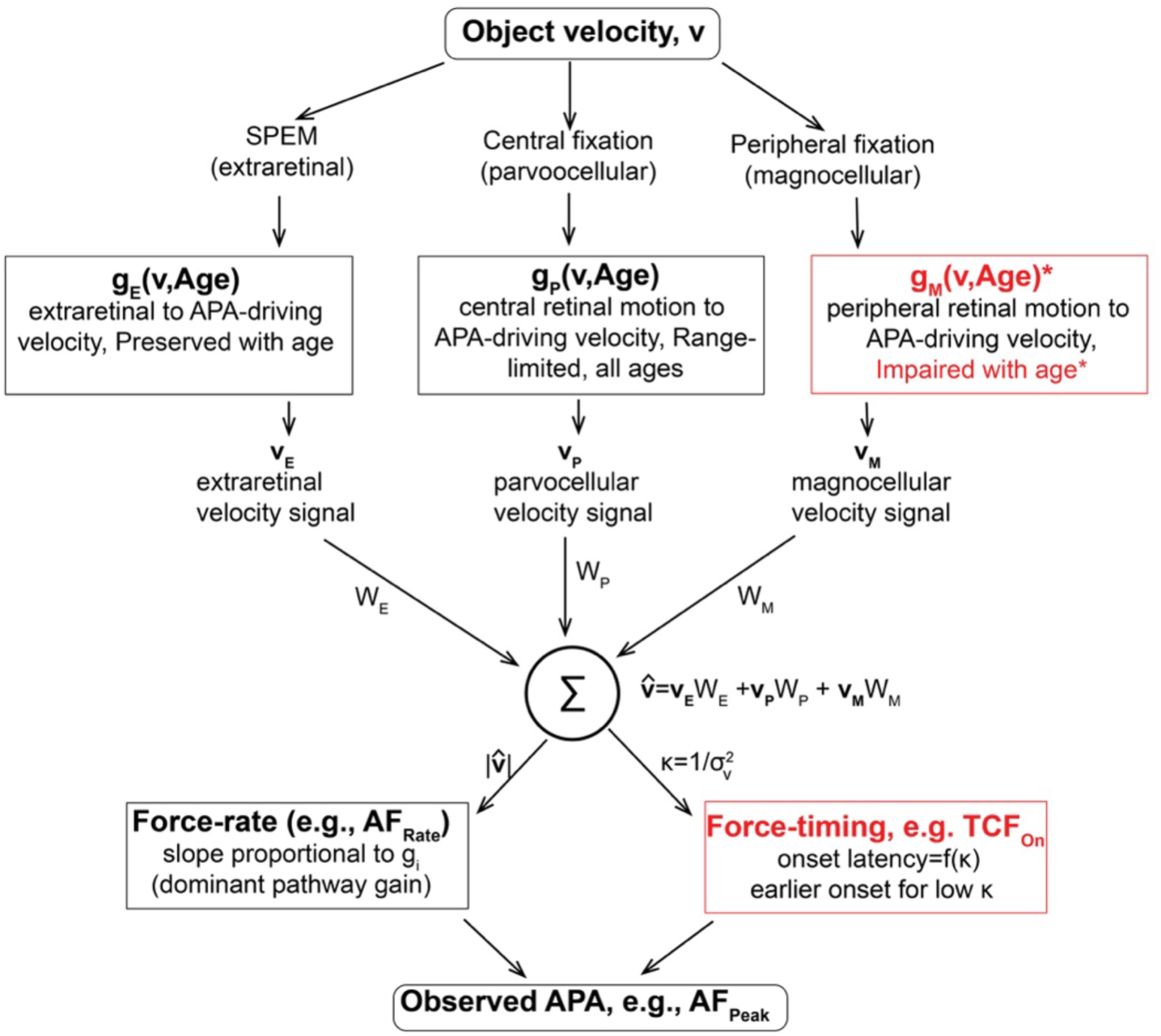
Conceptual model of aging effects on visually-driven anticipatory postural adjustments. Object velocity v is encoded in parallel by three motion-processing pathways selected by gaze condition: an extraretinal channel during smooth pursuit (SPEM), a magnocellular channel during peripheral fixation (FLR), and a parvocellular channel during central fixation (FC). Each pathway carries a gain function – g_E_(v, Age), g_M_ (v,Age), g_P_(v,Age) — defined as the input-to-APA coupling at velocity v in age group Age. Importantly, g_E_ refers to the coupling between the pursuit motor command and the APA-driving velocity signal and is distinct from oculomotor SPEM gain (eye velocity / target velocity), which declines with age while g_E_ is preserved. Pathway outputs v_E_, v_P_, v_M_ are combined at a precision-weighted summing junction with weights w_E_, w_P_, and w_M_ that are modulated by gaze condition: in each condition one weight dominates while the others contribute at reduced levels. The summing junction produces a velocity estimate v” characterized by both magnitude |v”| and precision κ=1/σ* where high κ corresponds to a reliable estimate and low κ to a noisy one (cue-combination convention). Force-rate generation reads the magnitude of |v”|, producing an APA force-rate slope (e.g., AF_Rate_) proportional to the dominant pathway’s gain g_i_. Force-timing reads the precision κ, producing an onset latency that decreases as κ decreases (e.g., TCF_On_). Together the two branches generate the observed APA (e.g., AF_Peak_). Red borders and asterisks mark sites where aging produces qualitative changes in the underlying mechanism: the magnocellular gain g_M_ is impaired with age (primary site), and the timing computation in Stage 4 is engaged differently in older adults, who shift force onset earlier when κ is reduced (when v” eventually becomes reliable, as during SPEM and FLR, but not when the pathway itself is the bottleneck, as during FC, where κ remains low regardless of integration time). Force-rate generation is itself unchanged with age; the lower AF_Rate_ slope at FLR in older adults reflects degraded g_M_ input rather than a deficit in force-rate machinery.

The architecture’s value, however, lies in the predictions it generates that the present data do not test. First, simultaneously engaging two pathways through deliberate stimulus geometry — for instance, having older adults pursue an offset cross while an object passes through peripheral retina — should produce APA slopes exceeding either single-pathway condition and partially rescue the FLR deficit, because the preserved g_E_ supplements the degraded g_M_. Second, the temporal compensation should carry a flexibility cost: older adults should be disproportionately vulnerable to mid-trajectory velocity perturbations, committing to the original *v*” rather than updating online. Third, individual variation in magnocellular function within the older cohort, assessed by independent measures, should specifically predict the FLR slope deficit and not the SPEM or FC slopes. Fourth, training that selectively targets the magnocellular pathway should restore some of the FLR deficit while leaving FC unchanged, whereas generic visual training should affect both equally or neither.

Together these predictions sharpen the model’s commitment: aging acts in two qualitatively different ways. The magnocellular gain is impaired in the strict sense, its input-to-output transformation degraded, while the timing computation is reorganized rather than impaired, engaged differently in older adults to recover what the gain deficit would otherwise lose. This dissociation between impairment and adaptive reorganization makes the model falsifiable, and provides a framework within which future findings can extend, refine, or overturn the present results.

## Conclusion

The present study demonstrates that the effect of aging on anticipatory postural control is pathway-specific: velocity-dependent APA scaling is preserved during pursuit tracking, limited by parvocellular system constraints during central fixation in both age groups, and selectively impaired during peripheral fixation in older adults, consistent with age-related decline in magnocellular-mediated motion processing. Older adults compensate for slower force dynamics by initiating anticipatory responses earlier, preserving peak force output. These findings establish a direct link between peripheral motion processing and anticipatory postural control, and identify the magnocellular-mediated peripheral motion processing pathway as a candidate target for interventions aimed at reducing collision-related falls in older adults.

## Acknowledgements

The authors thank Angus Muttee and Taylor Rosenquist for their help with data collection.

## Data availability

All data and analysis code supporting this study are openly available on the Open Science Framework at https://osf.io/uga2p/.

